# A stapled peptide inhibitor of MDM2 enables pharmacological activation of p53 in zebrafish

**DOI:** 10.64898/2026.03.26.714438

**Authors:** Sania Kheder, Martin Krkoška, Filip Mihalič, Kim Kobar, Zdeněk Andrysik, Lars Bräutigam, Susanne Lindström, Jason N Berman, David P Lane, Dilraj Lama, Pavitra Kannan

## Abstract

Measuring the activity of the tumor suppressor p53 in living systems is essential for understanding its dysregulation in cancer and other conditions, such as aging and diabetes. Zebrafish (*Danio rerio*) are a powerful vertebrate model that enable such studies, due to the evolutionary conservation of p53 structure and function. However, p53 activity in zebrafish has mainly been assessed using pharmacological methods that induce DNA damage or have off-target effects, making it difficult to isolate p53-specific responses from broader stress responses. Here, by using biophysical assays, molecular dynamics, and molecular assays, we show that sulanemadlin, a stapled peptide inhibitor of MDM2, binds to zebrafish Mdm2 and transcriptionally activates downstream targets of p53, including *cdkn1a*, isoform *Δ113p53*, and *Mdm2*. No effect on gene expression was observed in embryos treated with a point-modified control peptide or in embryos carrying a mutation that renders p53 transcriptionally inactive. RNA sequencing further confirmed upregulation of p53 signaling and downregulation of DNA replication pathways, while an acridine orange assay showed no detectable increases in apoptosis. In contrast, the tested small molecule Mdm2 inhibitors exhibit reduced binding affinity to zebrafish Mdm2 due to an amino acid variation in the zebrafish Mdm2 binding pocket. By overcoming a species-specific barrier in p53-MDM2 binding, the stapled peptide sulanemadlin is the first pharmacological tool to specifically activate p53 in zebrafish without inducing measurable apoptosis, enabling direct *in vivo* studies of p53 regulation in cancer and other disease contexts.

## INTRODUCTION

The p53 protein is an important transcription factor whose activation in response to distinct external stressors leads to different cellular outcomes. While it is rapidly degraded under unstressed conditions, p53 becomes stabilized under stress and activates downstream transcriptional targets that mediate cell cycle arrest, repair, senescence, and/or apoptosis^1^. Recent studies have shown that the transcriptional program activated by p53 is regulated not just by tissue type^2–4^ and the duration of activation^5^, but also by the nature of the external stress^6^. As a result, p53 activation leads to different cellular outcomes depending on context. Studying p53 transcriptional programs within relevant physiological or pathophysiological contexts is therefore essential for identifying how it regulates cellular stress response in conditions such as cancer, aging, and diabetes.

Zebrafish (*Danio rerio*) are a valuable model system for studying p53 activity because p53 structure and function are evolutionarily conserved. In terms of structure, the *tp53* gene in zebrafish is generally well-conserved, especially within key residues of the DNA-binding domain^7^. As a result, the canonical functional outcomes of p53, such as cell cycle arrest or apoptosis, are also conserved: induction of p53 increases the expression of downstream genes involved in cell cycle arrest and apoptosis^8^. However, due to its rapid degradation, p53 activity in zebrafish is generally induced using high dose radiation^9^ and/or the small molecule roscovitine, an inhibitor of cyclin-dependent kinase 9^10,11^. While both approaches strongly activate p53, they also induce apoptosis and/or have off-target effects that confound interpretation of p53-specific functions^9–11^. More selective approaches are needed to activate p53 without apoptosis or off-target effects, particularly to study conditions such as aging or diabetes.

A promising alternative for activating p53 pharmacologically is the use of compounds that block p53 from being targeted for proteasomal degradation by its negative regulator, Mdm2. These so-called Mdm2 inhibitors work by mimicking the three key residues in the transactivation domain (TAD) of p53 that binds to the hydrophobic Mdm2 binding pocket^12^. As a result, they block the interaction between Mdm2 and p53, preventing p53 degradation and thereby stabilizing its activity. We and others have shown that inhibitors of Mdm2 activate p53 in humans and mouse models. Importantly, these inhibitors are generally on-target and do not induce DNA damage^12–14^. Despite their increasing use for studying p53 activity in mammalian systems^12^, their ability to activate p53 in zebrafish remains poorly characterized. We found only one study reporting transient activation of p53 by a small molecule nutlin-3 in zebrafish^9^. Given that the zebrafish p53 TAD has a substitution in one of the three key residues involved in Mdm2 binding^15,16^, we reasoned that this difference in sequence could reduce the binding affinity of Mdm2 inhibitors to zebrafish and thereby their ability to activate p53.

To test this hypothesis, we used computational and experimental approaches to evaluate whether small molecule and stapled peptide inhibitors of Mdm2 have the potential to activate the p53 pathway in zebrafish. We selected two small molecule Mdm2 inhibitors, nutlin-3a and navtemadlin, that are widely used to activate p53 in mammalian models^13,14^. We also tested the stapled peptide sulanemadlin, a compound that mimics the *α*-helical conformation of the native structure of the p53 TAD and thereby binds a larger surface area of the Mdm2 binding pocket^12^. Using biophysical assays and molecular dynamics, as well as transcriptional and apoptosis assays, we show that the stapled peptide sulanemadlin binds with strong affinity to zebrafish Mdm2 and induces the transcriptional activity of p53 without measurable increases in apoptosis. In contrast, the small molecule inhibitors bind with weak affinity to zebrafish Mdm2, due to an amino acid substitution in the Mdm2 binding pocket. Sulanemadlin can therefore be used as a novel pharmacological tool to selectively activate p53 activity in zebrafish models.

## RESULTS

### Small molecule MDM2 inhibitors have reduced binding affinity to zebrafish Mdm2

We evaluated the binding affinities of small-molecule and stapled-peptide inhibitors to zebrafish Mdm2 using a fluorescence polarization assay. For clarity, we refer to the zebrafish protein as zMdm2 and the human ortholog as hMDM2. When discussing general mechanisms or inhibitors, we use ‘MDM2’ to denote the protein family across species. Previous studies have shown that while the native p53 protein has reduced affinity for zMdm2 compared to hMDM2^15^, the human p53 peptide binds to both zMDM2 and hMDM2 with similar affinity. Therefore, we first verified the binding affinity of human p53 peptide for the SWIB domains of both MDM2 proteins by measuring the displacement of fluorescein isothiocyanate (FITC) -labelled human p53 peptide (sequence: QETFSDLWKLL) by increasing concentrations of unlabeled human p53 peptide (SQETFSDLWKLLP) (**Figure 1A**). The human p53 peptide had affinity for both zMdm2 and hMDM2 (**Table 1**).

**Figure 1.**
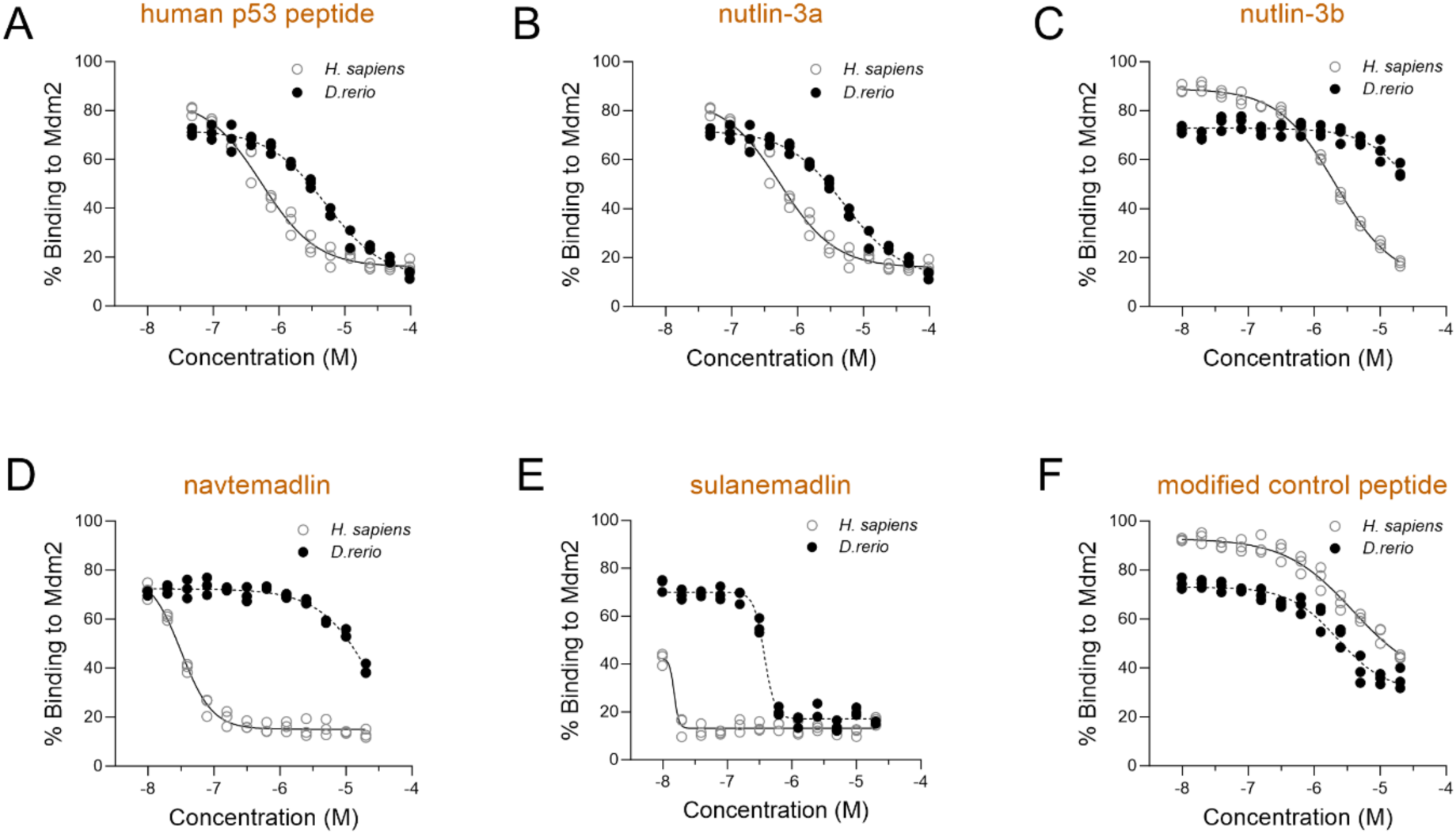
Small molecule MDM2 inhibitors are less effective than stapled peptides at displacing the binding of a p53 peptide to zebrafish Mdm2. The displacement of a fluorescently labelled human p53 peptide from human (MDM2, open circles) or zebrafish (Mdm2, filled circles) was measured using a fluorescence polarization assay at increasing concentrations of: **(A)** a human p53 peptide; **(B-D)** small molecules nutlin-3a, nutlin-3b (less potent enantiomer of nutlin-3a), and navtemadlin; **(E-F)** stapled peptide sulanemadlin and its modified control peptide (negative control). Each data point represents values from one independent experiment (n = 3 experiments). FP = fluorescence polarisation signal; mP = milli polarization units. Note that the EC_50_ values do not reflect the actual *K*_D_ value since they need to be recalculated based on specific experimental conditions as described in Nikolovska-Coleska et al^26^

**Table 1.** Binding affinities of small molecule and stapled peptide inhibitors of Mdm2/MDM2 to the zebrafish Mdm2 and human MDM2 proteins. The binding affinities were measured in vitro using a fluorescence polarization assay. * Due to the limitations of the assay, only the upper bound for affinity was estimated to be 10 nM. Real affinities are likely in sub-nanomolar range.

| Compound | Type | Zebrafish | Human |
| --- | --- | --- | --- |
| | | KD (μM) ± SD | KD (μM) ± SD |
| Human p53 peptide | | 0.943 ± 0.09 | 0.116 ± 0.02 |
| nutlin-3a | small molecule | 0.594 ± 0.07 | <0.01* ± n/a |
| nutlin-3b | small molecule | 8.463 ± 0.91 | 0.486 ± 0.01 |
| navtemadlin | small molecule | 3.094 ± 0.13 | <0.01* ± n/a |
| sulanemadiln | stapled peptide | <0.01* ± n/a | <0.01* ± n/a |
| modified control peptide | stapled peptide | 0.785 ± 0.11 | 1.946 ± 0.16 |

The ability of the inhibitors to displace the fluorescent FITC-p53 peptide from MDM2 was then measured using the same displacement assay. The small molecule nutlin-3a showed a nearly 60-fold weaker binding affinity to zMdm2 compared to hMDM2 (600 nM for zMdm2 vs. <10 nM for hMDM2; **Figure 1B**; **Table 1**). The less active enantiomer nutlin-3b, which was used as a negative control, had reduced affinity in both species (8.4 µM for zMdm2 and 486 nM for hMDM2; **Figure 1C**; **Table 1**). Although the small molecule navtemadlin is reported to be a more potent MDM2 inhibitor than nutlin-3a in mammalian systems^13,14,17–19^, it showed a ∼300-fold lower affinity for zMdm2 (3 µM for zMdm2 vs. <10 nM for hMDM2; **Figure 1D**; **Table 1**). Strikingly, its affinity in zebrafish was 5-fold lower than that of nutlin-3a. In contrast to the small molecules, the stapled peptide sulanemadlin had low nanomolar or subnanomolar affinity for MDM2 in both species (<10 nM; **Figure 1E**; **Table 1**). However, a modified version of the peptide containing a D-Phe substitution (henceforth called modified control peptide) had reduced binding in both species (780 nM for zMdm2 and 1.9 µM for hMDM2; **Figure 1F**; **Table 1**). Together, this biophysical assay shows that while the binding affinities of small molecule MDM2 inhibitors are reduced in zebrafish, the stapled peptide sulanemadlin retains exceptionally high affinity for both zMDM2 and hMDM2.

### A point substitution in the p53 binding pocket of zebrafish Mdm2/Mdm4 may reduce binding affinity of small molecules but not of the stapled peptide

To investigate the potential molecular basis underlying the reduced binding affinity of small molecule inhibitors in zebrafish compared to the stapled peptide, we performed molecular dynamics simulations of their modelled complexes with both zebrafish and human MDM2/MDM4. Analysis of the residue-wise binding energy decomposition from these simulations revealed that the major interactions of navtemadlin with MDM2/MDM4 in both species were localized in two regions (residues 52 to 62 and 92 to 102) of the proteins (**Figures 2A and 3A**). A key observation that emerges from a comparison of the energy profile is the contribution at residue position 96 in hMDM2, which shows a markedly higher binding energy (<-1kcal/mol) relative to the structurally equivalent position in zMdm2, as well as in MDM4 from both species (**Figures 2A and 3A**). In hMDM2, this position is occupied by histidine (H96), which forms a stabilizing hydrogen bond with navtemadlin in the bound state (**Figure 2B**). By contrast, zMdm2 and human/zebrafish MDM4 have a proline (P95/P92) substitution at this position, which results in the loss of hydrogen-bond interaction and a diminished binding energy contribution (**Figures 2B and 3B**). Interestingly, we have previously shown that navtemadlin efficiently reduces the growth of mouse melanoma cells in a p53-dependent manner^13^, consistent with effects observed in human cells^17^. Indeed, H96 is conserved in mouse Mdm2, and it forms a hydrogen bond with navtemadlin (**Supplementary Figure 1**). These results further support that the effectiveness of the compound could vary in a species-dependent manner. Importantly, H96 lies within the core of the p53 binding pocket, suggesting that this hydrogen bond could be a key molecular determinant for the higher binding affinity of navtemadlin for hMDM2 compared to zMdm2, or to MDM4 from either species.

**Figure 2.**
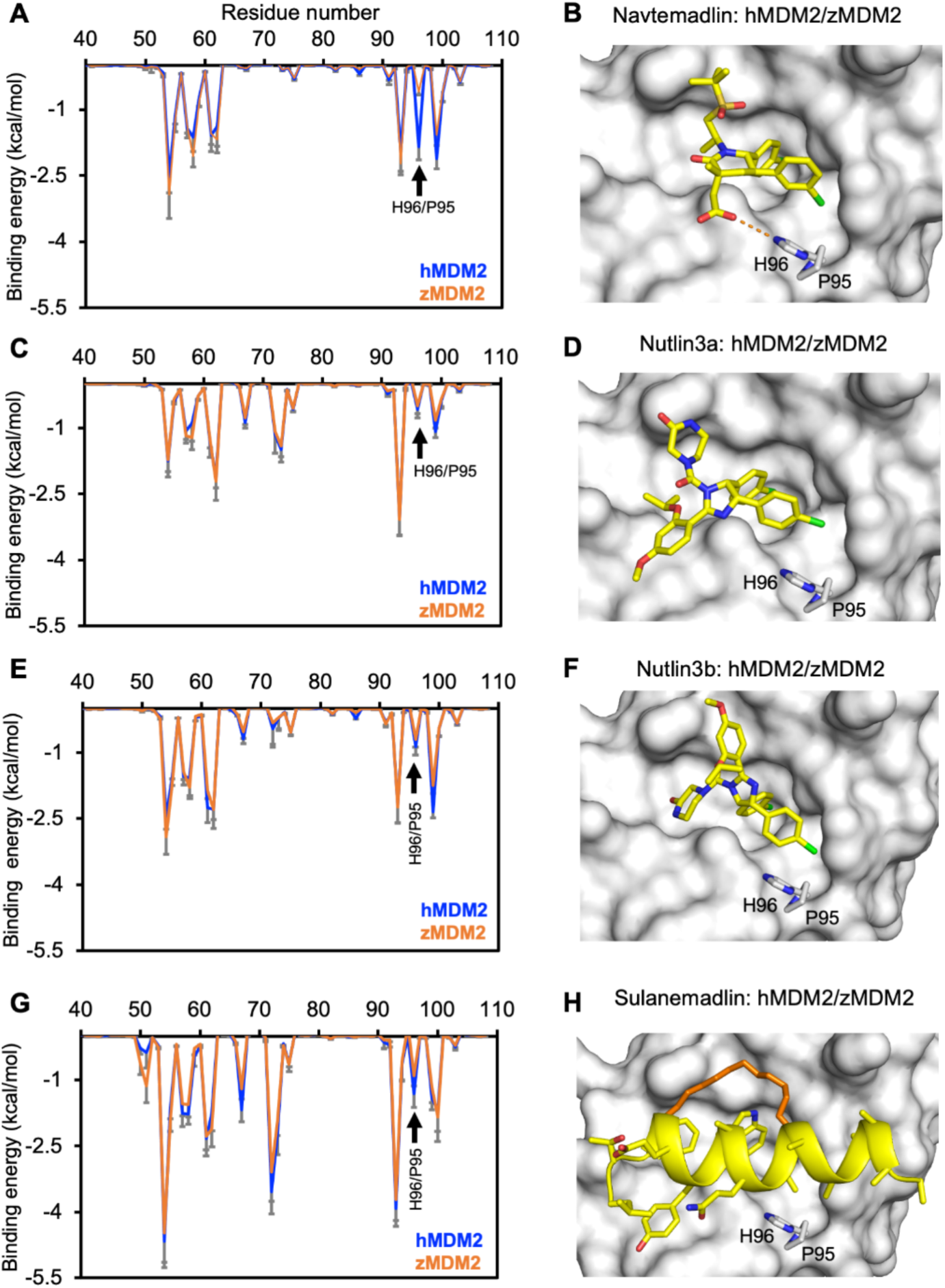
Energy decomposition of MDM2 complexes with small molecules and stapled peptide. **(A)** Residue-wise binding energy contribution (Ave±SD) from human and zebrafish MDM2 in complex with navtemadlin The H96/P95 residue position is indicated with an arrow. **(B)** Representative structure of human and zebrafish MDM2 in complex with navtemadlin. MDM2 is shown in surface, while navtemadlin and H96/P95 residues are depicted in stick representation. The hydrogen bond between H96 and navtemadlin is indicated with dotted lines. **(C)** Residue-wise binding energy contribution from human and zebrafish MDM2 in complex with nutlin-3a. **(D)** Representative structure of human and zebrafish MDM2 in complex with nutlin-3a. The figure is shown in similar representation except for the absence of the dotted lines indicating the lack of hydrogen-bond interaction in this system. **(E)** Residue-wise binding energy contribution from human and zebrafish MDM2 in complex with nutlin-3b. **(F)** Representative structure of human and zebrafish MDM2 in complex with nutlin3b. **(G)** Residue-wise binding energy contribution from human and zebrafish MDM2 in complex with sulanemadlin. **(H)** Representative structure of sulanemadlin in complex with human and zebrafish MDM2. The backbone and side chain of sulanemadlin is shown in cartoon and stick representation respectively. The staple linker is colored in orange.

**Figure 3.**
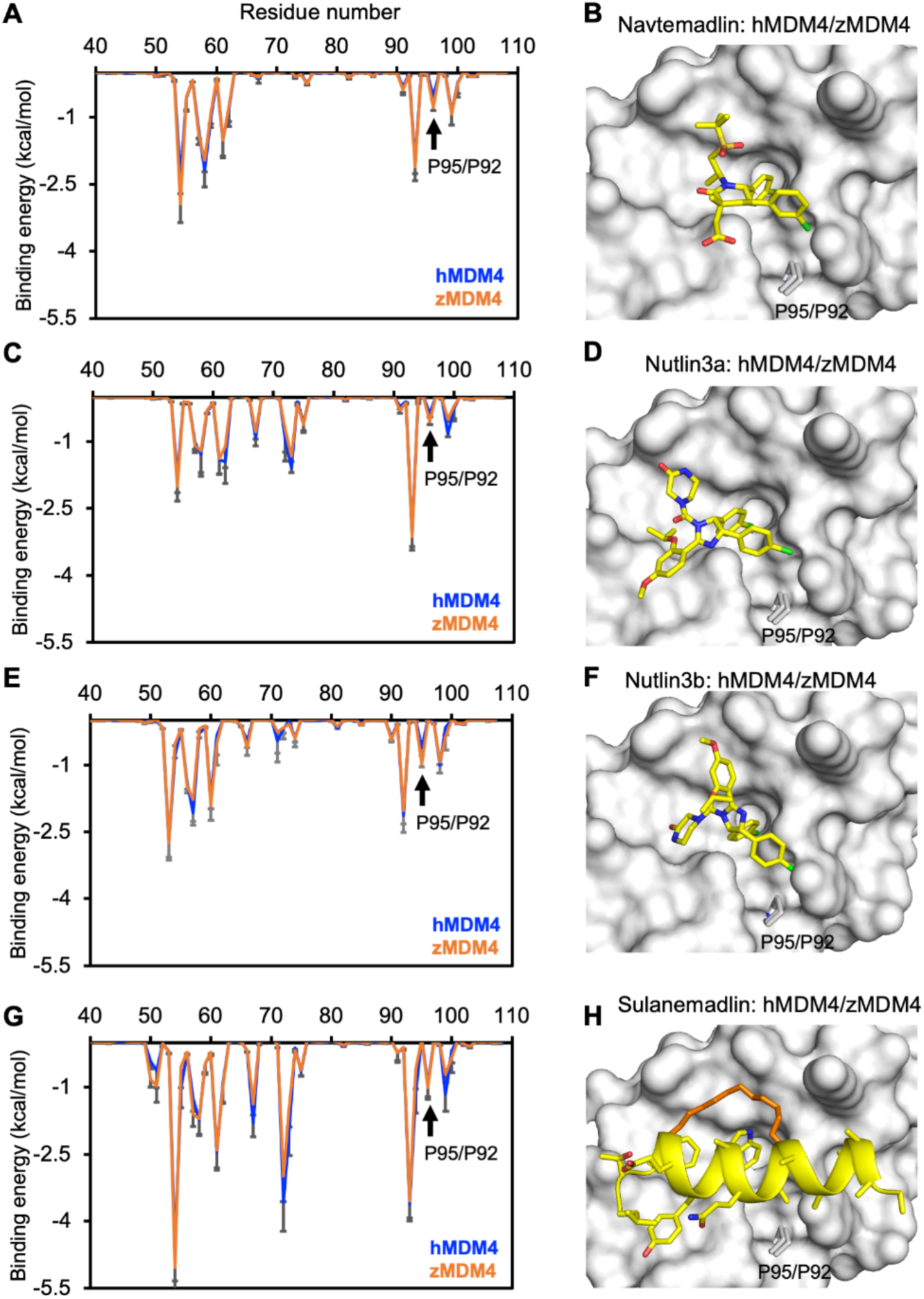
Energy decomposition of MDM4 complexes with small molecules and stapled peptide. **(A)** Residue-wise binding energy contribution (Ave±SD) from human and zebrafish MDM4. The P95/P92 residue position is indicated with an arrow. **(B)** Representative structure of human and zebrafish MDM4 in complex with navtemadlin. MDM4 is shown in surface, while navtemadlin and P95/P92 residues are depicted in stick representation. **(C)** Residue-wise binding energy contribution from human and zebrafish MDM4 in complex with nutlin-3a. **(D)** Representative structure of human and zebrafish MDM4 in complex with nutlin-3a. **(E)** Residue-wise binding energy contribution from human and zebrafish MDM4 in complex with nutlin-3b. **(F)** Representative structure of human and zebrafish MDM4 in complex with nutlin-3b. **(G)** Residue-wise binding energy contribution from human and zebrafish MDM4 in complex with sulanemadlin. **(H)** Representative structure of sulanemadlin in complex with human and zebrafish MDM4. The backbone and side chain of sulanemadlin is shown in cartoon and stick representation respectively. The staple linker is colored in orange.

We next performed the same analysis with complexes of nutlin-3a bound to MDM2/MDM4, which revealed that the major binding residues were largely similar to those for navtemadlin (**Figures 2C and 3C**). A notable difference, however, was observed at residue H96 in hMDM2. Its binding energy contribution was lower and comparable to that of zMdm2 and Mdm4 from both species. This reduction arises from the absence of a stabilizing hydrogen-bond interaction with H96 in the nutlin-3a: MDM2 complex (**Figures 2D and 3D**). The energy decomposition of the enantiomer nutlin-3b bound to MDM2/MDM4 showed a broadly similar binding energy profile as nutlin-3a (**Figures 2E and 3E**). However, due to differences in stereochemistry, the piperazinone fragment of nutlin-3b adopts a different spatial arrangement within the p53 binding pocket compared to nutlin-3a (**Figures 2F and 3F**). This altered orientation likely compromises critical protein-ligand interactions, which could explain the markedly reduced binding activity for the nutlin-3b enantiomer.

Finally, we examined the binding energy contributions in complexes of sulanemadlin with MDM2/MDM4. Compared to the small molecules, individual residues contributed relatively higher binding energies in this system (**Figures 2G and 3G**), though no stable hydrogen-bond interaction was observed with H96 in hMDM2 (**Figure 2H**). Sulanemadlin engages the p53-binding pocket of both human and zebrafish MDM2/MDM4 by mimicking the helical conformation, preserves the three native p53 interaction residues (F19, W23 and L26), and associates over a broader interaction interface than the small molecules (**Figures 2H and 3H**). In addition, the staple linker provides extra contacts with the protein, further strengthening the interaction. These features collectively account for the observed higher residue-wise binding energies and enable sulanemadlin to associate efficiently with MDM2/MDM4. As a result, the lack of the hydrogen-bond interaction with the histidine residue (or histidine to proline substitution) does not significantly influence sulanemadlin recognition of either human or zebrafish MDM2/MDM4. Taken together, these data suggest that the specific interaction with the histidine residue in the p53 binding pocket of MDM2 is a critical factor in defining the affinity for the small-molecule inhibitors, but not the stapled peptide.

### Stapled peptide sulanemadlin induces p53 transcriptional activity in zebrafish

Given that our biophysical and molecular dynamics assays indicated that stapled peptide sulanemadlin retains binding affinity for zMdm2, we subsequently measured its ability to transcriptionally activate seven key target genes in the p53 pathway using RT-qPCR (**Figure 4A**). Following 14 h treatment of embryos (48 hours post-fertilization) expressing wildtype p53, ≥1 μM sulanemadlin did not significantly change the expression of full-length *tp53* (*q-*value = 0.68). However, it significantly increased the expression of *mdm2* (negative regulator of p53; *q-*value _10μM_ = 0.004), *Δ113p53* (an isoform of p53 known that is transcriptionally activated by full length p53 in zebrafish; *q-*value _10μM_ = 0.004, *q-*value _1μM_ = 0.02), as well as the expression of *cdkn1a* (downstream target of cell cycle arrest; *q-*value _10μM_ = 0.001, *q-*value _1μM_ = 0.02; **Figure 4B**). However, sulanemadlin treatment did not increase the expression of any of the other genes, including that of negative regulator (*mdm4: q-*value _10μM_ = 0.99) and that of apoptosis genes (*baxa*: *q-*value _10μM_ = 0.99 and *puma*: *q-*value _10μM_ = 0.99; **Figure 4B**). The modified control peptide did not significantly increase the expression of any of the genes at any of the tested concentrations (ANOVA *F*_drug x gene_(18,47) = 1.023, *P* = 0.45), whereas the positive control roscovitine induced the expression of *Δ113p53* and *cdkn1a* (**Figure 4B**), consistent with its reported activity in zebrafish^20^.

**Figure 4.**
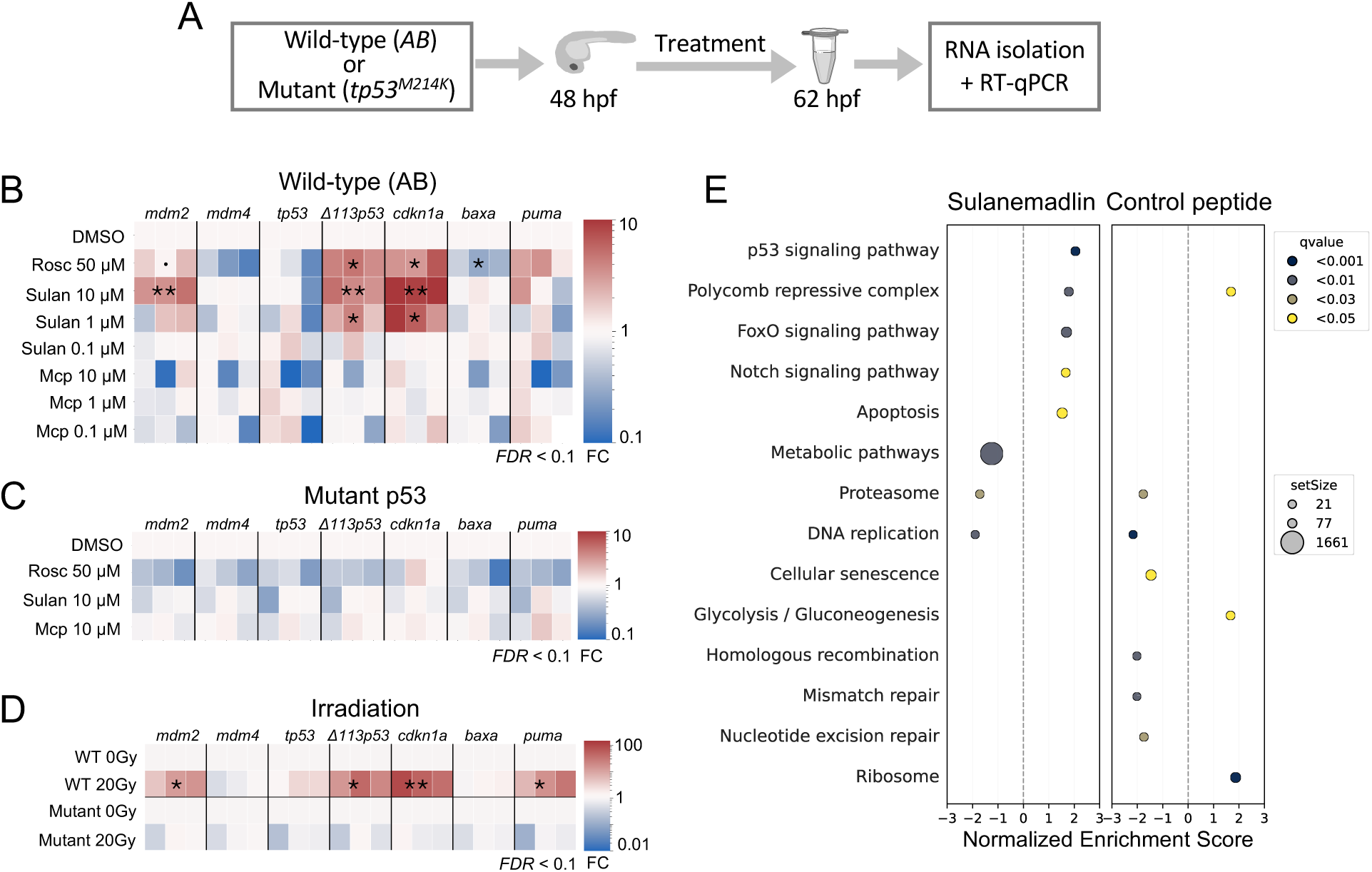
Stapled peptide MDM2 inhibitor sulanemadlin induces p53 transcriptional activity in zebrafish embryos. **(A)** Schematic showing workflow for zebrafish embryo treatments and collection of lysates. **(B)** Increased expression of p53 downstream targets, *mdm2*, *Δ113tp53,* and *cdkn1a*, following ≥ 1 μM treatment with sulanemadlin (sulan) or 50 μM roscovitine (rosc) but not with increasing concentrations of the modified control peptide (Mcp). **(C)** No change in expression is measured in zebrafish expressing a point mutation in p53 at M214K^9^. **(D)** Increased expression of several p53 downstream targets (*mdm2*, *Δ113tp53*, *cdkn1a*, and *puma*), following treatment radiation of 20 Gy. Statistical significance was evaluated using ANOVA followed by false discovery rate (FDR) of 0.1 using the Benjamini, Krieger, and Yekutieli. **(E)** Gene set enrichment analysis of KEGG pathways shows upregulation of p53 signaling by sulanemadlin but not by the modified control peptide. Normalized enrichment scores are shown for significantly enriched pathways (q-value < 0.05). The dot sizes reflect the number of genes found in the ranked expression list, with circle sizes indicating the minimum, median, and maximum values.

To confirm that the measured transcriptional responses required functional p53, we then measured the expression of the same genes in zebrafish embryos homozygous for M214K point mutation in the DNA binding domain of p53 (tp53^zdf1/zdf1^), which renders p53 transcriptionally inactive (**Figure 4C**). None of the tested compounds (i.e., roscovitine, sulanemadlin, and control peptide) had a significant effect on the expression of any of the tested genes in the mutant M214K zebrafish line (ANOVA *F*_drug x gene_(18,48) = 1.348, *P* = 0.20). Finally, to rule out that the lack of apoptosis gene expression induced by the compounds was due to technical limitations of the RT-qPCR, we measured the expression of *puma* and *baxa* in wildtype AB and mutant p53 zebrafish embryos treated with 20 Gy radiation (**Figure 4C**), which is known to induce apoptosis^9^. In irradiated wildtype AB fish, *puma* increased > 2-fold (*q-*value = 0.04), but *baxa* did not increase significantly (*q-*value = 0.21). In contrast, no significant changes were measured in any of the genes from irradiated mutant p53 embryos (ANOVA *F*_radiation dose x gene_(6,24) = 0.4753, *P* = 0.82).

To identify other molecular targets that might be affected by sulanemadlin treatment in zebrafish embryos, we subsequently performed bulk RNA sequencing on lysates from wildtype zebrafish embryos treated with DMSO, 50 μM roscovitine, 10 μM sulanemadlin, and 10 μM control peptide. Given that principal component analysis indicated high inter-replicate variability (which reduces statistical power to detect a large number of differentially expressed genes), we performed gene set enrichment analysis of KEGG pathways to identify biological processes that were most affected with each treatment. Following sulanemadlin treatment, the p53 signaling, Notch signaling, and apoptosis pathways were upregulated, whereas DNA replication was among the downregulated ones (**Figure 4E**). In contrast, treatment with control peptide did not show an increase in the p53 signaling pathway, but did show an effect on the DNA replication pathway. In contrast, roscovitine induced substantial changes in many functional and biological pathways, including upregulation of ribosome, oxidative phosphorylation, glycolysis, p53 signaling pathways, and downregulation of nucleotide excision repair and metabolism (**Supplementary Figure 2**). To further validate the qPCR results, we confirmed that *mdm2* (log2FC = 2.29; *P_raw_* = 4.24×10^-6^) and *ckdn1a* (log2FC = 2.13; *P_raw_* = 0.003) were both increased in RNA seq data, as analyzed by edgeR. Taken together, these results demonstrate that the stapled peptide sulanemadlin induces p53 transcriptional activity in a p53-dependent manner and is on-target.

### Sulanemadlin does not measurably induce apoptosis in zebrafish embryos

Based on the results of the RNA sequencing, we subsequently tested whether treatment led to a phenotypic increase in the number of apoptotic cells in zebrafish embryos. Therefore, we used an acridine orange assay to detect apoptotic cells (**Supplementary Figure 3**) following 6 h compound treatment in 30 hpf embryos, a developmental stage during which apoptosis is detectable in multiple tissues^21^. In wild-type fish, none of the test compounds (sulanemadlin, the modified control peptide, or roscovitine), increased the number of apoptotic cells compared to DMSO (**Figures 5A, 5B**). As expected, the positive control camptothecin, which induces apoptosis through p53^22^, induced a ∼2.7-fold increase in apoptotic cells (*P_adj_*< 0.001; **Figures 5A, 5B**). In either p53-null embryos or embryos carrying the p53^R242H/R242H^ mutation, no significant increase in apoptotic cells above baseline levels was measured for any of the compounds (**Figures 5A, 5B**). Together, these findings confirm that although sulanemadlin may transcriptionally activate genes within the apoptotic pathway, it does not induce a measurable increase in apoptotic cells within the experimental timeframe tested here.

**Figure 5.**
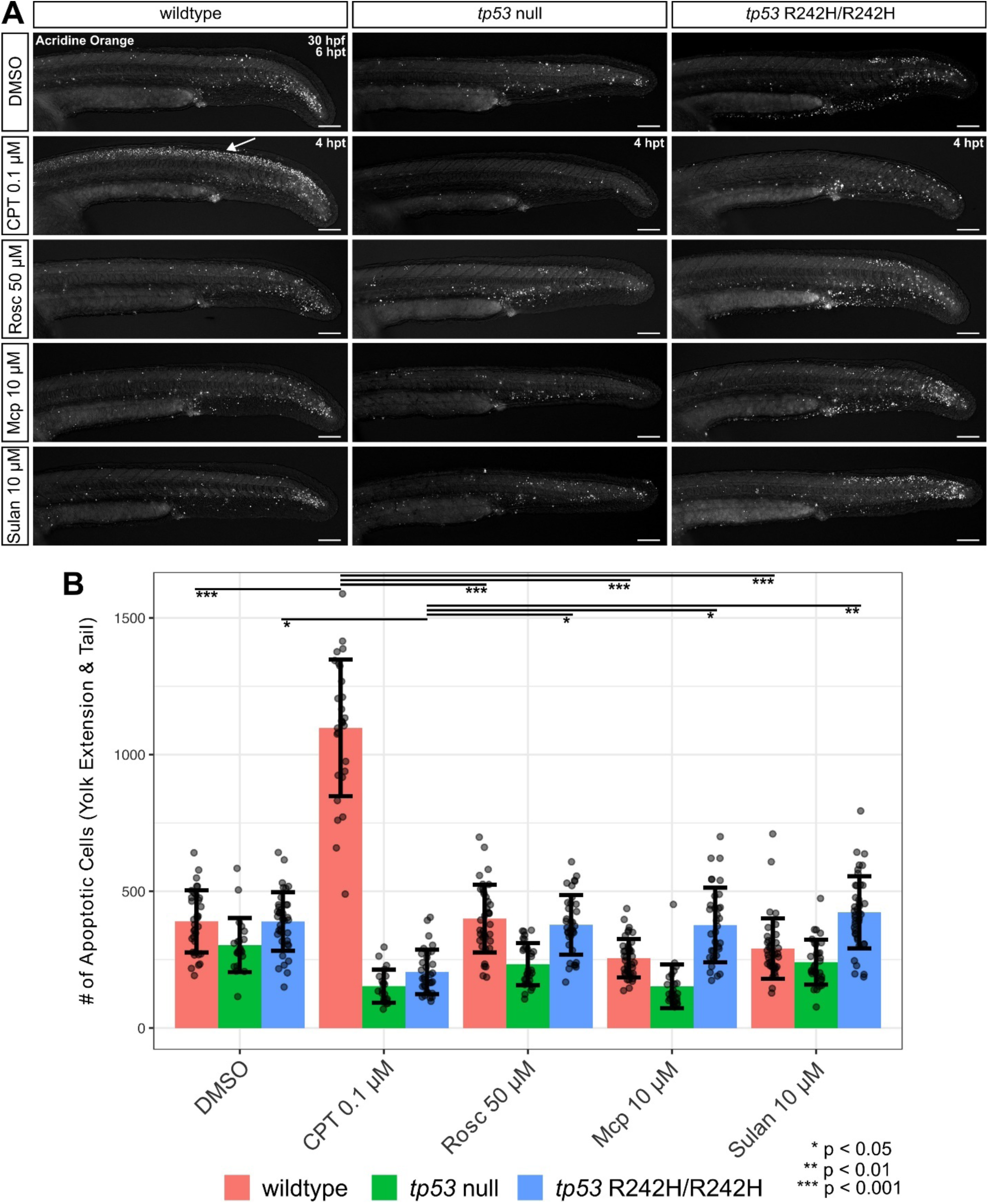
Sulanemadlin treatment does not measurably increase apoptosis in wildtype, *tp53* null, or *tp53^R242H/R242H^* zebrafish embryos. **(A)** Representative images of the yolk extension and tail of acridine orange stained 30 hpf wildtype, *tp53^fb105/fb105^* null, and *tp53^R242H/R242H^* zebrafish embryos treated with DMSO, camptothecin (CPT), roscovitine (rosc), modified control peptide (Mcp), or sulanemadlin (sulan) at 24 hpf (CPT treatment was at 26 hpf for 4 hours). The white arrow highlights the typical strong apoptosis pattern along the dorsal spine. Scale bars, 500μm. **(B)** Quantification of the number of apoptotic cells in the tail and area above the yolk extension. Error bars represent standard deviation. Each data point represents one embryo pooled from three independent biological replicates. Outliers were identified and excluded using Grubbs’ test with alpha < 0.05. Statistical significance was determined by a two-way ANOVA with correction for multiple testing using a Tukey’s honest significance difference test, as indicated by asterisks.

## DISCUSSION

### Species differences in binding of MDM2 inhibitors

Using a combination of biophysical, computational, and biological experiments, we evaluated the efficacy of small molecule and stapled peptide inhibitors of MDM2 in zebrafish. We found that while small molecule MDM2 inhibitors have weaker binding affinity to zMdm2 than to hMDM2, stapled peptide sulanemadlin retains binding affinity and activates p53 transcriptional activity *in vivo* without a measurable increase in apoptotic cells. These results show that the stapled peptide sulanemadlin is a novel and specific pharmacological tool to transcriptionally activate p53 in zebrafish embryos.

A key result in our study is that the binding affinity of small molecule MDM2 inhibitors is weaker for zMdm2 than for hMDM2, as evidenced by our biophysical and computational analysis. This reduced affinity may be explained by the key differences in amino acid substitution of histidine to proline in zebrafish within the p53 binding pocket of zMdm2. Since the histidine residue is in the core of the p53 binding pocket, the hydrogen-bond interaction between the compound and the binding pocket likely contributes towards the high affinity of navtemadlin in humans and in mice, but not in zebrafish. The hydrogen-bond interaction between navtemadlin and the binding pocket could also explain why this compound has a higher affinity than nutlin-3a to hMDM2, as well as why its affinity is substantially affected in zebrafish as compared to nutlin-3a. In contrast, the conservation of helical scaffold, native mode of binding, and the additional interactions of the stapled peptide can promote an efficient association of MDM2 ^12^. Consequently, the lack of the hydrogen-bond interaction with the histidine residue is not a significant factor in its recognition of either zebrafish or human MDM2. Thus, the specific interaction with the histidine residue in the p53 binding pocket is likely a critical factor in defining the affinity for small molecule but not the stapled peptide.

### Transcriptional activity induced by Mdm2 inhibition

We found that the stapled peptide induced p53 transcriptional activity in zebrafish embryos, as measured by RT-qPCR assays of downstream transcriptional targets and functional pathway analysis from whole genome sequencing. At ≥ 1 μM, the stapled peptide significantly induced the expression of cell cycle arrest gene *cdkn1a* by nearly 10-fold, but not that of canonical apoptosis genes *baxa* or *puma*. Functional pathway analysis partly corroborated this data, showing that the top upregulated pathway by 10 μM sulanemadlin was p53 signaling. Although apoptosis also emerged as an enriched pathway, we did not observe any overt functional apoptosis at the tested concentrations in zebrafish embryos using an in vivo acridine orange assay. These data are in line with previous results: we and others have previously shown that stapled peptide MDM2 inhibitors on their own primarily induce cell cycle arrest and only moderate levels of apoptosis in mouse and human cells^13,14,23^. Another explanation for the limited apoptotic response is that the isoform *Δ113tp53* reduces the apoptosis activity of full-length *tp53* in zebrafish^20^. We found that the stapled peptide significantly induced the expression of the isoform by nearly 10-fold at 14 h after treatment, similar to the effects induced by roscovitine in previous studies^9,10,20^. It is possible that the activity of full-length *tp53* is induced at earlier time points and then reduced following the induction of the isoform, as has been reported before. Given that the molecular structure of *Δ113tp53* lacks the zMdm2-interacting motif^20^, it is unlikely that the isoform binds the stapled peptide but is rather transcriptionally activated by full-length *tp53* itself. Future studies could be conducted in *Δ113tp53*^-/-^ embryos to determine whether apoptosis activity is still induced by the stapled peptide.

We also investigated how the activation pattern of the stapled peptide compares to that induced by roscovitine and radiation. We found that the stapled peptide was more potent than roscovitine at inducing similar expression patterns in 48 hpf zebrafish embryos: 1 μM sulanemadlin induced nearly the same level of *cdkn1a* and *Δ113tp53* activation as 50 μM roscovitine. While both compounds induced cell cycle arrest by about 10-fold, the levels were much lower than that induced by radiation, which induced arrest and apoptosis by 30-fold or more. These results are consistent with recent reports showing that different transcriptional activation patterns are induced by different stressors^6,24^. It is unclear whether the induction of gene expression could be increased further through MDM2 inhibition in zebrafish embryos, but we did not test this possibility. Thus, the selective activation of cell cycle arrest over apoptosis genes by sulanemadlin represents a substantial advantage for studying p53 activity in developmental or aging models of zebrafish, where apoptotic activity may be harmful to normal tissues and confound interpretation of results.

### Limitations of the study

While our study demonstrates that the stapled peptide can activate p53 in zebrafish, several aspects require further exploration. First, we did not consider tissue-specific differences in activation patterns^2^, which could be addressed in future studies using single cell RNA sequencing or transgenic reporter lines. Second, as with any pharmacological tool, a characterization of the biodistribution of the stapled peptide within different zebrafish tissues is needed. Third, our study provides only a snapshot of p53 activation at a single time point following treatment, rather than capturing the temporal dynamics of p53 activity following treatment. Although previous studies have shown that p53 activation starts around 2 h and peaks at around 14 h following roscovitine treatment^9,20^, the duration of activity for the stapled peptide remains unclear. Finally, affinities below 10 nM are impossible to accurately measure with our fluorescence polarization assay. While the upper limits for the affinities of nutlin-3a, navtemadlin, and sulanemadlin for hMDM2 and sulanemadlin for zMdm2 were estimated to be 10 nM, the true affinities are likely in the sub-nanomolar range, as reported previously^19,25^.

## CONCLUSION

In summary, using a combination of biophysical, computational, and biological techniques, we demonstrate that the stapled peptide sulanemadlin binds with high affinity to zMdm2 and induces the cell cycle arrest activity of p53 without inducing apoptosis. This selective activation represents an advantage to conventional methods that induce apoptosis or have off-target effects. Since sulanemadlin does not measurably induce apoptosis in zebrafish, it can be used as a novel tool to study the context-specific activity of p53 in zebrafish models of aging, cancer, and diabetes.

## AUTHOR CONTRIBUTIONS

PK conceived the study with input from DL and DPL. Data were acquired by SK, MK, FM, KK, ZA, LB, SL, and DL. Data analysis and interpretation were performed by SK, MK, FM, KK, ZA, DPL, DL, and PK. PK wrote the initial manuscript draft with input from DL, FM, KK, ZA, and DPL. All authors contributed to the revision and editing of the manuscript. Project supervision was performed by JNB, DPL, DL, and PK, while funding was provided by ZA, JNB, DPL, and PK.

## ACKNOWLEDGEMENTS

This project has received funding from the European Union Horizon 2020 research and innovation programme under the Marie Sklodowska-Curie grant agreement No. 839585 (PK). The research was also supported by the Swedish Research Council (DPL; 2013-08807 and 2024-03430), the Gunvor and Josef Anérs Foundation (PK; FB23-0164), the Lillian Sagens and Curt Ericssons Research Foundation (PK; 4-591/2025), the Swedish Cancer Society (PK; 22 0527 FE), and by the Czech Science Foundation project No. 25-18368S (ZA).

This work is dedicated to the memory of Professor Per Jemth, who provided resources and funding for the biophysical assays. Professor Jemth passed away before the manuscript was completed. We thank Heidi Josefsson for technical assistance and the zebrafish core facility (Karolinska Institutet) for support and assistance with experiments. We acknowledge the Central European Institute of Technology (Masaryk University) CF Genomics and the CF Bioinformatics supported by the NCMG research infrastructure (LM2023067 funded by MEYS CR) for their help with obtaining RNA-seq data presented in this paper. Part of the schematic in Fig. 4A was made using graphics from Servier Medical Art, licensed under a Creative Commons Attribution 3.0 Unported License (https://smart.servier.com/).

## DATA AVAILABILITY

The datasets generated during the current study are available in the Zenodo repository (DOI: web link to be made available). Any other relevant data are available upon reasonable request from the corresponding authors.

## MATERIALS AND METHODS

### In vitro experiments

#### Chemicals

We tested three MDM2 inhibitors in zebrafish embryos: the small molecules nutlin-3a and navtemadlin, and the stapled peptide sulanemadlin. A modified control peptide was used as a negative control. The small molecule roscovitine, an inhibitor of cyclin dependent kinase 9, was used as a positive control to induce p53 activity in the embryos. Camptothecin (MilliporeSigma, C9911) was used as a positive control to induce apoptosis in zebrafish embryos. Sulanemadlin (Ac-Leu-Thr-Phe-R8-Glu-Tyr-Trp-Ala- Gln-Leu-S5-Ala-Ala-Ala-Ala-Ala-DAla- NH2; olefin staple from R8 to S5) and the modified control peptide (Ac-Leu-Thr-dPhe-R8-Glu-Tyr-Trp-Ala- Gln-Leu-S5-Ala-Ala-Ala-Ala-Ala-DAla- NH2; olefin staple from R8 to S5) were custom synthesized (BioSynth). Compounds were dissolved in DMSO, aliquoted, and stored at −80 °C at the following concentrations: 10 mM nutlin-3a, 10 mM nutlin-3b, 10 mM navtemadlin, 10 mM sulanemadlin, 10 mM control peptide, and 50 mM roscovitine. For assays, they were dissolved to the final concentration in E3 medium. Camptothecin was stored as 2 mM stock in DMSO at −20 °C and dissolved to final concentration in methylene-blue free E3 medium. All compounds were purchased from MedChemExpress, unless specified otherwise.

#### MDM2 protein purification and binding affinity measurements using fluorescence polarization

Human and zebrafish MDM2 protein purifications and binding affinity determinations were performed as described previously^16^. In short, the *E. coli* BL21 (DE3) strain was transformed with pETM33 plasmids coding for either human or zebrafish MDM2 SWIB domains, grown in 2xYT medium at 37 °C untill OD_600_ reached 0.6-0.8 and the protein expression was induced with the addition of IPTG (1 mM final concentration). After overnight expression at 18 °C cells were harvested and resuspended in lysis buffer (50 mM Tris/HCl pH 7.8, 300 mM NaCl, 10-µg/mL DNase I and RNase A, 4 mM MgCl_2_, 2 mM CaCl_2_, and cOmplete EDTA-free Protease Inhibitor Cocktail). Cell lysis was achieved with sonication and cell debris were palleted by centrifugation (20 000xg, 30 min). Lysates were applied to Pierce Glutathione Agarose beads (Thermo Scientific), washed with buffer (50 mM Tris, 300 mM NaCl, and pH 7.8), and eluted with 10 mM reduced glutathione. The GST tag was cleaved overnight at 4 °C using PreScission protease and the cleavage reaction was applied to Nickel Sepharose Fast Flow column (GE Healthcare) to remove the GST tag. Final round of size exclusion chromatography was employed (column Hi load 16/60 Sephacryl S-100 column (GE Healthcare)) to remove any impurities and the final purified protein samples were dialyzed into experimental buffer (20 mM sodium phosphate, pH 7.4, 150 mM NaCl, and 1 mM TCEP) before being frozen at –80 until further use.

For fluorescence polarization affinity measurements, a previously established protocol was employed^16^. A fixed concentration of fluorescein isothiocyanate (FITC) labeled human p53 peptide (15 nM) (QETFSDLWKLL) was premixed with either human or zebrafish MDM2 (final concentration of 100 nM and 750 nM respectively) and added to an increasing concentration of displacer peptide/molecule (two fold dilution series; highest concentrations of displacer compounds were 96.7 µM for human p53 peptide, 26 µM for control peptide and 20 µM for all other compounds; peptide sequences for human p53 displacer peptide was SQETFSDLWKLLP (Gencust)). Experiments were performed in experimental buffer at room temperature on SpectraMax iD5 plate reader (Molecular Devices), and the results were analysed using GraphPad Prism 9. Sigmoidal dose response (variable slope) fitting model was employed to obtain EC_50_ values and those were converted to *K*_D_ values according to Nikolovska-Coleska and colleagues^26^.

### In silico experiments

#### Molecular dynamics of MDM2/MDM4 with small molecules and stapled peptide

The structures of the p53-binding domain from human,mouse and zebrafish MDM2/MDM4 (Uniprot IDs: Q00987/O15151/P23804 and O42354/Q7ZUW7/O35618) respectively was downloaded from the AlphaFold database^27^. The coordinates of navtemadlin, nutlin-3a, nutlin-3b were obtained from Pubchem. Molecular docking of these ligands with MDM2 and MDM4 structures was done using Swissdock^28^. The stapled peptide sulanemadlin bound to MDM2 and MDM4 was derived by first modelling the linear peptide in complex with the protein using AlphaFold2 (AF2) through the jupyter notebook for ColabFold (version 1.3.0)^29^. The MSA_mode was set to MMseqs2 (UniRef+Environment), pair_mode selected as “unpaired+paired”, model type employed was “AlphaFold-multimer-v2” with three number of recycles, no template information was used, and amber relaxation of models was disabled. The i, i+7 hydrocarbon linker (staple), whose force-field parameters were derived as previously described^30^, was then modelled into the structure. The terminal ends of the protein domain was capped with ACE and NME functional groups. The structures were separately placed in the centre of a truncated octahedral box, whose dimensions were fixed by setting at-least 10 Å between any protein atom and the box edges. TIP3P water model^31^ was used for solvation, and the net charge neutrality of the individual system was ensured by adding appropriate number of counter ions. MD simulations were carried out using the PMEMD module of AMBER18^32^ employing ff19SB force-field parameters^33^. The systems were energy minimized, heated to 300 K over 30 ps and equilibrated for 200 ps under NVT and NPT ensembles respectively. Finally, the production run was executed for 100 ns each for all the systems under the NPT conditions. A harmonic positional restrain of 5 kcal/mol was applied to all the backbone heavy atoms of the protein/stapled-peptides and the small molecules. Langevin dynamics^34^ (with collision frequency of 1.0 ps^-1^) and weak coupling^35^ (with relaxation time of 1 ps) was used to maintain simulation temperature of 300 K and pressure of 1 atm respectively. Periodic boundary conditions were applied and the Particle Mesh Ewald^36^ was used to compute long-range electrostatic interactions. All the hydrogen containing bonds was constrained with the SHAKE^37^ algorithm, and the equation of motion was solved with an intergration time step of 4 fs by applying hydrogen-mass repartitioning^38^.

#### Residue-wise binding energy decomposition

Residue-wise free energy decomposition was calculated with the Molecular Mechanics / Generalized Born Surface Area (MM/GBSA) method^39^. A total of 1000 structures covering the simulation period were extracted at equal intervals. Explicit water molecules and counter ions were stripped from the extracted structures, and the solvent effect was represented using an implicit Generalized Born solvation model (IGB=2). The dielectric constant of the solvent was set to 80, ***γ***=0.0072 kcal/mol/Å^2^ and salt concentration to 150 mM. The bonded and non-bonded energies of the protein was calculated using Molecular Mechanics force field expression of AMBER, the polar solvation energy was obtained with the GB solvation model, and the non-polar solvation energies was estimated using an empirical linear relationships (***γ***\*SASA), where ***γ*** is the surface tension and SASA is the solvent accessible surface area. The MMPBSA.py^40^ script available through the AMBER18 suite of programs was used for the computation.

### In vivo experiments

#### Animal husbandry and ethics

Two genetic strains of zebrafish embryos (*Danio rerio*) were used to measure the transcriptional activity induced by pharmacological p53 activation. The wildtype strain (AB) and p53 mutant strain expressing a point mutation at M214K (*tp53*^zdf1/zdf1^)^41^ were housed at Karolinska Institutet, as previously described^42^. The p53 mutant strain was purchased from European Zebrafish Resource Center (Germany). Zebrafish embryos were staged according to published guidelines^43^. At Karolinska Institutet, animals were monitored for health status by Charles River according to FELASA-AALAS guidelines; Mycobacterium chelonae and Mycobacterium fortuitum were found in randomly sampled fish and sludge samples as previously reported. Animal procedures were approved by the Stockholm Regional Animal Ethics Committee (Dnr 15591/2023). The procedures were performed in compliance with relevant guidelines and regulations under the Swedish Board of Agriculture’s Regulations and General Advice of Laboratory Animals (Statens jordbruksverks föreskrifter och allmänna råd om försoksdjur; SJVFS 2019:9; Saknr L150) and EU legislation (Directive 2010/63/EU).

Three genetic strains were used to measure apoptotic activity induced by pharmacological p53 activation: wildtype (*casper* AB), *tp53^fb105/fb105^* null on CG1 background, and *tp53* ^R242H/R242H^ on *casper* AB background. Zebrafish were maintained at 28 °C on a recirculating water system with a 14:10 h light:dark cycle at the University of Ottawa Aquatics Facility. Embryos were staged as per published guidelines^43^. Animal use and procedures were approved by and conducted in accordance with the University of Ottawa’s Animal Care Committee under protocol #4166, which is in line with the Canadian Council for Animal Care’s guidelines.

#### Pharmacological treatment of zebrafish embryos

For transcriptional assays, embryos (48 h post-fertilization, hpf) were dechorionized and incubated for 14 h at 28.5 °C in E3 medium containing 160 µg/mL tricaine, 30 µg/mL phenylthiourea, and either solvent (DMSO 0.1%) or one of three compounds (50 μM roscovitine, 10, 1 or 0.1 μM sulanemadlin, or 10, 1 or 0.1 μM modified control peptide). Embryos were then collected for downstream processing.

For acridine orange apoptosis assays, dechorionated embryos (24 hpf) were placed in 12-well plates and incubated for 6 h at 28 °C in E3 medium containing one of four compounds (50 μM roscovitine, 10 μM sulanemadlin, 10 μM modified control peptide, or DMSO). For 0.1 μM camptothecin, embryos were treated at 26 hpf for 4 h at 28 °C and washed three times with methylene blue-free E3 before acridine orange staining.

#### Irradiation of zebrafish embryos

To verify the ability to measure apoptosis in our assays embryos (48 hpf) were irradiated using X-rays (X-Strahl) at a dose rate of 2.39 Gy/min (Al filter of 3 mm, FSD = 30 cm). Embryos were collected 5 hours later for downstream processing.

#### Transcriptional activity using RT-qPCR

To detect whether the stapled peptides and roscovitine activate p53 transcriptional activity in zebrafish, we performed RT-qPCR. Total RNA was extracted from embryo lysates collected in TRIzol™ Reagent (700 µL per 10 embryos; ThermoFisher # MAN0001271) using the Direct-zol™ RNA Miniprep Kit (Zymo Research #R2050), with DNAse I treatment. cDNA synthesis (700-1000 ng per reaction) was performed using the iScript cDNA Synthesis Kit (BIO-RAD #1708890), according to manufacturer’s instructions. For RT-qPCR, eight genes (Table 2) were selected: the full length *tp53* and its isoform *delta113 tp53*, the downstream targets *cdkn1a*, *mdm2*, and *mdm4*, the p53-dependent regulators of apoptosis *bax* and *puma*, and the housekeeping gene *b2m*. The efficiency of the primers (Integrated DNA Technologies) was confirmed to be within the range of 90–110% using the standard curve method. Each qPCR reaction was performed in a 96-well plate (Applied Biosystems #434907) using 500 nM primer mix, 2.5 ng cDNA, and PowerTrack™ SYBR Green Master Mix (ThermoFisher # A46012). The data were analyzed using StepOnePlus™ Software v2.3 and normalized using the ΔΔCt method. Heatmaps of the fold changes were created in Python (v. 3.8.8) using the pandas, numpy, matplotlib and seaborn libraries, with coding assistance from Claude Sonnet (4.0). The outputs were manually verified.

**Table 2.**
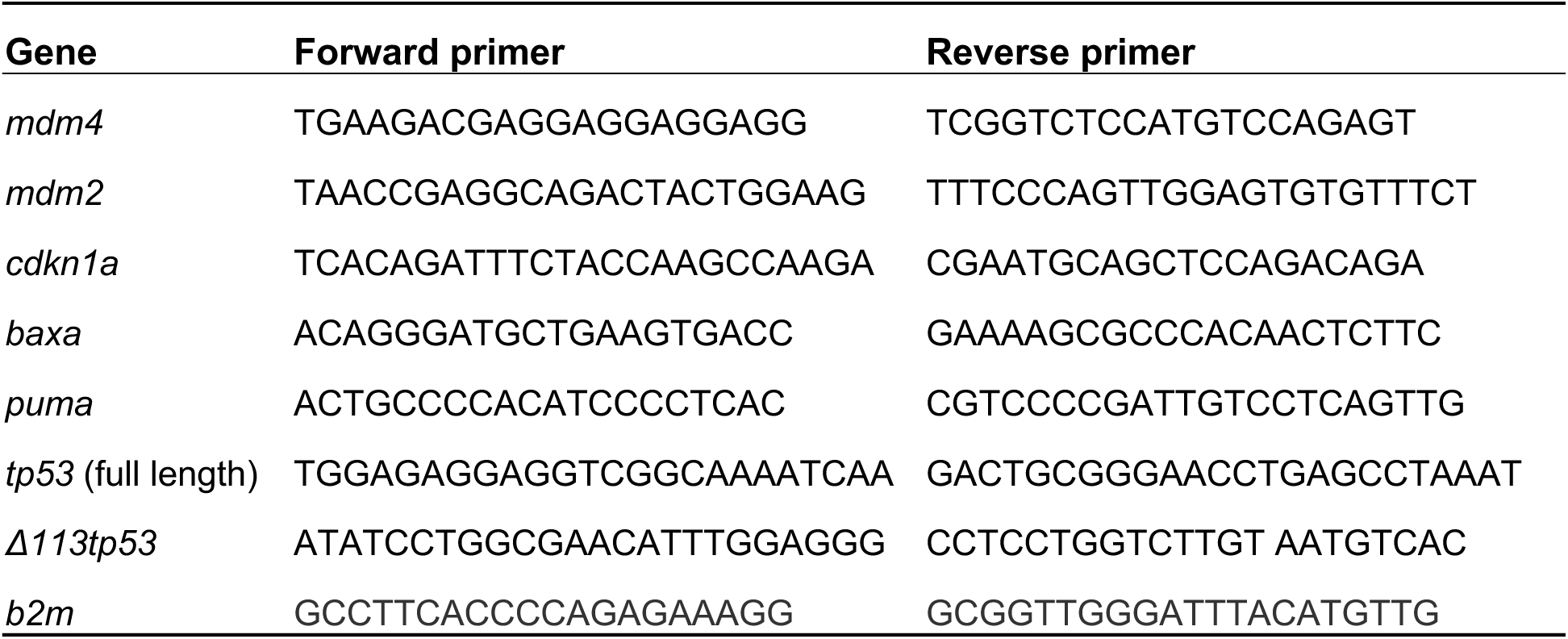
Primer sequences used for quantitative PCR to measure gene expression in zebrafish embryo lysates.

#### RNA sequencing and gene set enrichment analysis of KEGG pathways

To identify whole genome changes induced by sulanemadlin treatment, we performed RNA sequencing on zebrafish embryos treated with DMSO, 50 μM roscovitine, 10 μM sulanemadlin, or 10 μM of the modified control peptide. The RNA extracted for RT-qPCR was also used for sequencing. RNA integrity was assessed using TapeStation High Sensitivity RNA ScreenTape Assay (Agilent). Independent biological triplicates were used to prepare sequencing libraries (QuantSeq, Lexogen), which were processed at AVITI24 sequencer (Element Biosciences) resulting in >11 million reads per sample. After adapter trimmed, single-end reads were quality filtered and aligned to *Danio rerio* reference genome using STAR aligner (GRCz11, Ensembl r113). Differential gene expression was analyzed using EdgeR. Gene set enrichment analysis (GSEA) of KEGG pathways was performed on ranked expression data DESeq2 v1.34.0. Normalized enrichment scores of significantly enriched pathways were visualized as dot plots in Python (v. 3.8.8) using the pandas, numpy, and matplotlib libraries, with coding assistance from Claude Sonnet (4.0). Outputs were manually verified.

#### Acridine Orange Staining, Imaging, and Quantification

To assess whether sulanemadlin induced apoptosis, we performed an acridine orange assay. Dechorionated and drug-treated embryos (30 hpf) were immersed in 2 μg/mL acridine orange (Thermo Fisher, A1301) for 30 minutes and washed 4x with methylene blue-free E3 medium. Embryos were anesthetized with 160 μg/mL tricaine and placed into 96 well plates, 1 embryo/well. Embryos were imaged using the FITC (GFP/FITC/Cy2 Filter Set) and RGB brightfield channels. Images were captured using an in-house automated pipeline on a Nikon TI2-E inverted microscope equipped with a Digital Sight 10 CMOS camera (Nikon Instruments Inc.). Image visualization and processing was done in Fiji and NIS-elements AR 6.02.00 software. Segment.ai (NIS-Elements AR software) was used to segment the yolk extension and tail region (Supplemental Figure 3) for each image, and all binaries were checked manually to ensure proper segmentation. The number of apoptotic cells in the yolk extension and tail were counted using a General Analysis 3 task (NIS-Elements AR software) to sharpen the image and apply the bright spot detection analysis in the Segment.ai binary region.

### Statistical analysis

The biophysical experiments were performed as three independent experiments, with the averaged values from each experiment reported as mean ± standard deviation. The RT-qPCR experiments were performed as three biological experiments, each comprising 10-12 pooled embryos. For each drug, the effect of treatment on gene expression was measured using one-way or two-way ANOVA (GraphPad Prism v.10) with factors drug and gene. If a drug x gene interaction was identified as significant, a subsequent post-hoc test was performed using the Benjamini, Krieger, and Yekutieli at a false discovery rate of 0.1. The model fit was assessed using residuals and Q-Q normality plots. Zebrafish samples were excluded if they failed quality control (e.g., poor RNA quality). Randomization and blinding were not performed.

## REFERENCES

1 Boutelle AM, Attardi LD. p53 and Tumor Suppression: It Takes a Network. Trends Cell Biol 2021; 31: 298–310.

2 Pant V, Sun C, Lozano G. Tissue specificity and spatio-temporal dynamics of the p53 transcriptional program. Cell Death Differ 2023; 30: 897–905.

3 Moyer SM, Wasylishen AR, Qi Y, Fowlkes N, Su X, Lozano G. P53 drives a transcriptional program that elicits a non-cell-autonomous response and alters cell state in vivo. PNAS 2020; 117: 23663–23673.

4 Bowen ME, McClendon J, Long HK, Sorayya A, Van Nostrand JL, Wysocka J et al. The Spatiotemporal Pattern and Intensity of p53 Activation Dictates Phenotypic Diversity in p53-Driven Developmental Syndromes. Dev Cell 2019; 50: 212–228.e6.

5 Hafner A, Bulyk ML, Jambhekar A, Lahav G. The multiple mechanisms that regulate p53 activity and cell fate. Nat Rev Mol Cell Biol 2019; 20: 199–210.

6 Zhou RW, Silverman L, Manfredi JJ, Parsons RE, 4. Basal p53 maintains a distinct transcriptional program from irradiated p53 in tissue, including tumor suppressors. bioRx 2024; 15: 37–48.

7 Kobar K, Collett K, Prykhozhij S V., Berman JN. Zebrafish Cancer Predisposition Models. Front Cell Dev Biol 2021; 9: 1–27.

8 Mandriani B, Castellana S, Rinaldi C, Manzoni M, Venuto S, Rodriguez-Aznar E et al. Identification of p53-target genes in Danio rerio. Sci Rep 2016; 6: 1–13.

9 Guo L, Liew HP, Camus S, Goh AM, Chee LL, Lunny DP et al. Ionizing radiation induces a dramatic persistence of p53 protein accumulation and DNA damage signaling in mutant p53 zebrafish. Oncogene 2013; 32: 4009–4016.

10 Lee KC, Goh WLP, Xu M, Kua N, Lunny D, Wong JS et al. Detection of the p53 response in zebrafish embryos using new monoclonal antibodies. Oncogene 2008; 27: 629–640.

11 Jerafi-Vider A, Bassi I, Moshe N, Tevet Y, Hen G, Splittstoesser D et al. VEGFC/FLT4-induced cell-cycle arrest mediates sprouting and differentiation of venous and lymphatic endothelial cells. Cell Rep 2021; 35: 109255.

12 Tuval A, Strandgren C, Heldin A, Palomar-Siles M, Wiman KG. Pharmacological reactivation of p53 in the era of precision anticancer medicine. Nat Rev Clin Oncol 2024; 21: 106–120.

13 Ingelshed K, Spiegelberg D, Kannan P, Påvénius L, Hacheney J, Jiang L et al. The MDM2 Inhibitor Navtemadlin Arrests Mouse Melanoma Growth In Vivo and Potentiates Radiotherapy. Cancer Res Commun 2022; 2: 1075–1088.

14 Clavero AL, Boqvist PL, Ingelshed K, Bosdotter C, Sedimbi S, Jiang L et al. MDM2 inhibitors, nutlin-3a and navtemadelin, retain efficacy in human and mouse cancer cells cultured in hypoxia. Sci Rep 2023; 13: 1–12.

15 Mihalič F, Åberg E, Farkhondehkish P, Theys N, Andersson E, Jemth P. Evolution of affinity between p53 transactivation domain and MDM2 across the animal kingdom demonstrates high plasticity of motif-mediated interactions. Protein Sci 2023; 32: 1–17.

16 Mihalič F, Arcila D, Pettersson ME, Farkhondehkish P, Andersson E, Andersson L et al. Conservation of Affinity Rather Than Sequence Underlies a Dynamic Evolution of the Motif-Mediated p53/MDM2 Interaction in Ray-Finned Fishes. Mol Biol Evol 2024; 41: 1–13.

17 Canon J, Osgood T, Olson SH, Saiki AY, Robertson R, Yu D et al. The MDM2 inhibitor AMG 232 demonstrates robust antitumor efficacy and potentiates the activity of p53-inducing cytotoxic agents. Mol Cancer Ther 2015; 14: 649–658.

18 Vassilev L, Vu BT, Graves B, Carvajal D, Podlaski F, Filipovic Z et al. In Vivo Activation of the p53 Pathway by Small-Molecule Antagonists of MDM2. Science (80-) 2004; 303: 844–849.

19 Sun D, Li Z, Rew Y, Gribble M, Bartberger MD, Beck HP et al. Discovery of AMG 232, a Potent, Selective, and Orally Bioavailable MDM2–p53 Inhibitor in Clinical Development. J Med Chem 2014; 57: 1454–1472.

20 Guo L, Chua J, Vijayakumar D, Lee KC, Lim K, Eng H et al. Detection of the 113p53 protein isoform: A p53-induced protein that feeds back on the p53 pathway to modulate the p53 response in zebrafish. Cell Cycle 2010; 9: 1998–2007.

21 Kobar K, Tuzi L, Fiene JA, Burnley E, Galpin KJC, Midgen C et al. tp53 R217H and R242H mutant zebrafish exhibit dysfunctional p53 hallmarks and recapitulate Li-Fraumeni syndrome phenotypes. Biochim Biophys Acta - Mol Basis Dis 2025; 1871: 167612.

22 Rudolf E, Cervinka M, Rudolf K. Camptothecin induces p53-dependent and -independent apoptogenic signaling in melanoma cells. Apoptosis 2011; 16: 1165–1176.

23 Ingelshed K, Melssen MM, Kannan P, Chandramohan A, Partridge AW, Jiang L, et al. MDM2/MDMX inhibition by Sulanemadlin synergizes with anti-Programmed Death 1 immunotherapy in wild-type p53 tumors. iScience 2024; 27: 109862.

24 Stewart-Ornstein J, Iwamoto Y, Miller MA, Prytyskach MA, Ferretti S, Holzer P et al. P53 Dynamics Vary Between Tissues and Are Linked With Radiation Sensitivity. Nat Commun 2021; 12. doi:10.1038/s41467-021-21145-z.

25 Carvajal LA, Ben Neriah D, Senecal A, Benard L, Thiruthuvanathan V, Yatsenko T et al. Dual inhibition of MDMX and MDM2 as a therapeutic strategy in leukemia. Sci Transl Med 2018; 10. doi:10.1126/scitranslmed.aao3003.

26 Nikolovska-Coleska Z, Wang R, Fang X, Pan H, Tomita Y, Li P et al. Development and optimization of a binding assay for the XIAP BIR3 domain using fluorescence polarization. Anal Biochem 2004; 332: 261–273.

27 Varadi M, Bertoni D, Magana P, Paramval U, Pidruchna I, Radhakrishnan M et al. AlphaFold Protein Structure Database in 2024: providing structure coverage for over 214 million protein sequences. Nucleic Acids Res 2024; 52: D368–D375.

28 Grosdidier A, Zoete V, Michielin O. SwissDock, a protein-small molecule docking web service based on EADock DSS. Nucleic Acids Res 2011; 39: 270–277.

29 Mirdita M, Schütze K, Moriwaki Y, Heo L, Ovchinnikov S, Steinegger M. ColabFold: making protein folding accessible to all. Nat Methods 2022; 19: 679–682.

30 Lama D, Liberatore AM, Frosi Y, Nakhle J, Tsomaia N, Bashir T et al. Structural insights reveal a recognition feature for tailoring hydrocarbon stapled-peptides against the eukaryotic translation initiation factor 4E protein. Chem Sci 2019; 10: 2489–2500.

31 Jorgensen WL, Chandrasekhar J, Madura JD, Impey RW, Klein ML. Comparison of simple potential functions for simulating liquid water. J Chem Phys 1983; 79: 926–935.

32 Case DA, Walker RC, Cheatham TE, Simmerling C, Roitberg A, Merz KM et al. Amber 18. Univ California, San Fr 2018 2018.http://ambermd.org/doc12/Amber18.pdf.

33 Tian C, Kasavajhala K, Belfon KAA, Raguette L, Huang H, Migues AN et al. Ff19SB: Amino-Acid-Specific Protein Backbone Parameters Trained against Quantum Mechanics Energy Surfaces in Solution. J Chem Theory Comput 2020; 16: 528–552.

34 Loncharich RJ, Brooks BR, Pastor RW. Langevin dynamics of peptides: The frictional dependence of isomerization rates of N-acetylalanyl-N′-methylamide. Biopolymers 1992; 32: 523–535.

35 Berendsen HJC, Postma JPM, Gunsteren WF van, DiNola A, Haak JR. Molecular dynamics with coupling to an external bath. J Chem Phys 1984; 81: 3684–3690.

36 Darden T, York D, Pedersen L. Particle mesh Ewald: An N⋅log(N) method for Ewald sums in large systems. J Chem Phys 1993; 98: 10089–10092.

37 Ryckaert J-P, Ciccotti G, Berendsen HJ. Numerical integration of the cartesian equations of motion of a system with constraints: molecular dynamics of n-alkanes. J Comput Phys 1977; 23: 327–341.

38 Hopkins CW, Le Grand S, Walker RC, Roitberg AE. Long-time-step molecular dynamics through hydrogen mass repartitioning. J Chem Theory Comput 2015; 11: 1864–1874.

39 Genheden S, Ryde U. The MM/PBSA and MM/GBSA methods to estimate ligand-binding affinities. Expert Opin Drug Discov 2015; 10: 449–461.

40 Miller BR, Mcgee TD, Swails JM, Homeyer N, Gohlke H, Roitberg AE. MMPBSA py : An Efficient Program for End-State Free Energy Calculations. J Chem Theory Comput 2012; 8: 3314–3321.

41 Berghmans S, Murphey RD, Wienholds E, Neuberg D, Kutok JL, Fletcher CDM et al. tp53 mutant zebrafish develop malignant peripheral nerve sheath tumors. PNAS 2005; 102: 407–412.

42 Al-Radi O, Ingelshed K, Eichhorn L, Josefsson H, Krkoska M, Bräutigam L et al. Pharmacological activation of p53 induces dose-dependent changes in endothelial cell fate during angiogenic sprouting. Cell Death Dis 2025; 16. doi:10.1038/s41419-025-08292-7.

43 Kimmel CB, Ballard WW, Kimmel SR, Ullmann B, Schilling TF. Stages of embryonic development of the zebrafish. Dev Dyn 1995; 203: 253–310.

