## Supplementary Information for "A stapled peptide inhibitor of MDM2 enables pharmacological activation of p53 in zebrafish"

**Running title:** Stapled peptide as a novel tool to study zebrafish p53

[#]**Equal contribution as first authors**

[†]**Equal contribution as last authors**

***Corresponding authors:**

Dilraj Lama,

Pavitra Kannan,

**FIGURES**

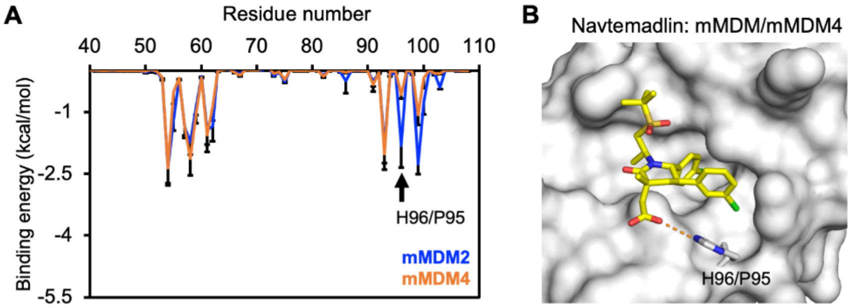


**Supplementary Figure 1. Energy decomposition of mouse MDM2 and MDM4 complexes with small molecules and stapled peptide. (A)** Residue-wise binding energy contribution (Ave±SD) from mouse MDM42 and MDM4. The H95/P96 residue position is indicated with an arrow. **(B)** Representative structure of mouse MDM2 and MDM4 in complex with navtemadlin. MDM2 and MDM4 is shown in surface, while navtemadlin and H95/P96 residues are depicted in stick representation.

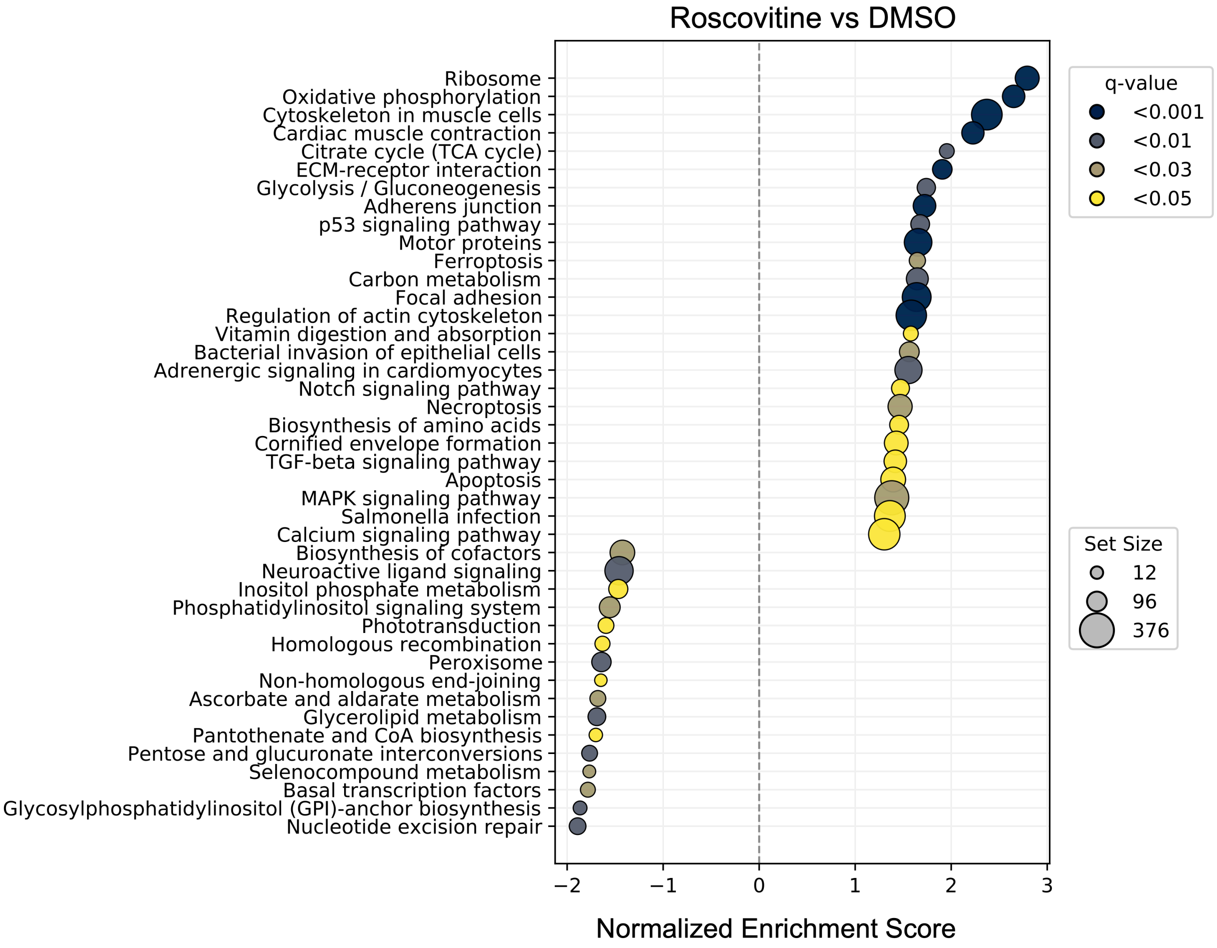


**Supplementary Figure 2. Small molecule roscovitine affects multiple functional pathways in zebrafish embryos.** Gene set enrichment analysis of KEGG pathways was performed on data from edgeR analysis. The normalized enrichment scores are shown for significantly enriched pathways (q-value < 0.05). The dot sizes reflect the number of genes found in the ranked expression list, with circle sizes indicating the minimum, median, and maximum values.

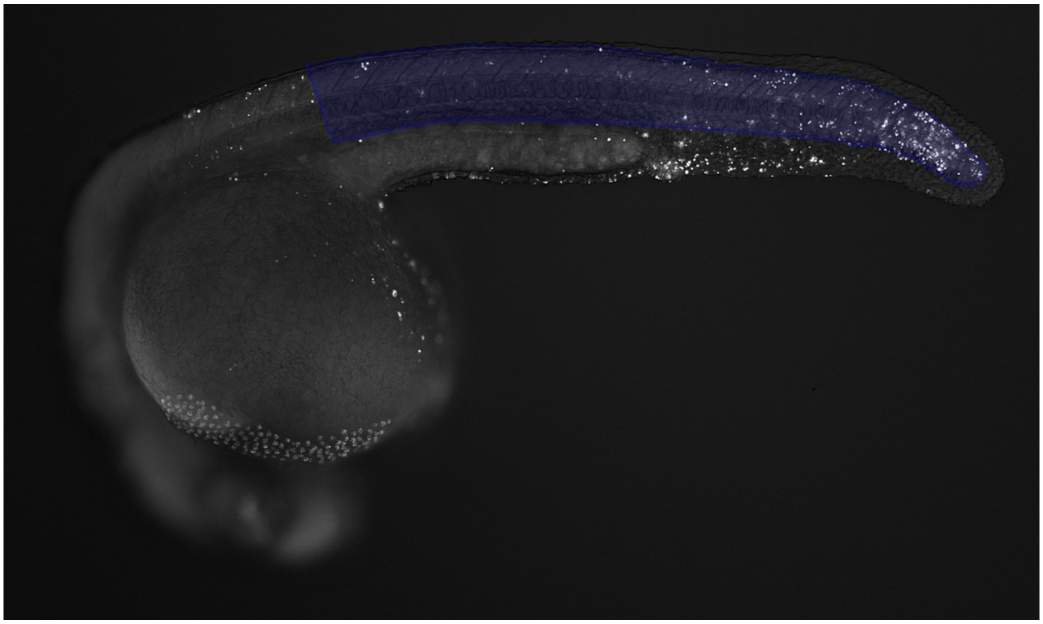

**Supplementary Figure 3. Representative image showing the tail and yolk extension area used for the apoptosis quantification.** Binary (blue) overlay highlights the area above the yolk extension and tail used for acridine orange apoptosis quantification. Binary generated using NIS-Elements AR Segment.ai (Nikon Instruments Inc.).
